# Intra-Species Differences in Population Size shape Life History and Genome Evolution

**DOI:** 10.1101/852368

**Authors:** David Willemsen, Rongfeng Cui, Martin Reichard, Dario Riccardo Valenzano

## Abstract

The evolutionary forces shaping life history trait divergence within species are largely unknown. Killifish (oviparous Cyprinodontiformes) evolved an annual life cycle as an exceptional adaptation to life in arid savannah environments characterized by seasonal water availability. The turquoise killifish (*Nothobranchius furzeri*) is the shortest-lived vertebrate known to science and displays differences in lifespan among wild populations, representing an ideal natural experiment in the evolution and diversification of life history. Here, by combining genome sequencing and population genetics, we investigate the evolutionary forces shaping lifespan among turquoise killifish populations. We generate an improved reference assembly for the turquoise killifish genome, trace the evolutionary origin of the sex chromosome, and identify genes under strong positive and purifying selection, as well as those evolving neutrally. We find that the shortest-lived turquoise killifish populations, which dwell in fragmented and isolated habitats at the outer margin of the geographical range of the species, are characterized by small effective population size and accumulate throughout the genome several small to large-effect deleterious mutations due to genetic drift. The genes most affected by drift in the shortest-lived turquoise killifish populations are involved in the WNT signalling pathway, neurodegenerative disorders, cancer and the mTOR pathway. As the populations under stronger genetic drift are the shortest-lived ones, we propose that limited population size due to habitat fragmentation and repeated population bottlenecks, by causing the genome-wide accumulation of deleterious mutations, cumulatively contribute to the short adult lifespan in turquoise killifish populations.

## Main

The extent to which drift and selection shape life history trait evolution across species in nature is a fundamental question in evolutionary biology. Variations in population size among natural populations is expected to affect the rate of accumulation of advantageous and slightly deleterious gene variants, hence impacting the relative contribution of selection and drift to genetic polymorphisms^1^. Populations living in fragmented habitats, subjected to continuous and severe bottlenecks, are expected to undergo dramatic population size reduction and drift, which can significantly impact the accumulation of genetic polymorphisms in genes affecting important life history traits^2^.

Among vertebrates, killifish represent a unique system, as they repeatedly and independently colonised highly fragmented habitats, characterized by cycles of rainfalls and drought^3^. While on the one hand intermittent precipitation and periodic drought pose strong selective pressures leading to the evolution of embryonic diapause, an adaptation that enables killifish to survive in absence of water^4,5^, on the other hand they cause habitat and population fragmentation, promoting inbreeding and genetic drift. The co-occurrence of strong selective pressure for early-life and extensive drift characterizes life history evolution in African annual killifishes ^6^.

The turquoise killifish (*Nothobranchius furzeri*) is the shortest-lived vertebrate with a thoroughly documented post-embryonic life, which, in the shortest-lived strains, amounts to four months^4,5,7,8^. Turquoise killifish has recently emerged as a powerful new laboratory model to study experimental biology of aging due to its short lifespan and to its wide range of aging-related changes, which include neoplasias^9^, decreased regenerative capacity^10^, cellular senescence^11,12^, and loss of microbial diversity^13^. At the same time, while sharing physiological adaptations that enable embryonic diapause and rapid sexual maturation, different wild turquoise killifish populations display differences in lifespan, both in the wild and in captivity^14-16^, making this species an ideal evolutionary model to study the genetic basis underlying life history trait divergence within species.

Characterisation of life history traits in wild-derived laboratory strains of turquoise killifish revealed that while different populations have similar rates of sexual maturation^8^, populations from arid regions exhibit the shortest lifespans, while populations from more semi-arid regions exhibit longer lifespans^8,14^. Hence, speed of sexual maturation and adult lifespan appear to be independent in turquoise killifish populations. The evolutionary mechanisms responsible for the lifespan differences among turquoise killifish populations are not yet clearly understood. Mapping genetic loci associated with lifespan differences among turquoise killifish populations showed that adult survival has a complex genetic architecture^15,17^. Here, combining genome sequencing and population genetics, we investigate to what extent genomic divergence in natural turquoise killifish populations that differ in lifespan is driven by adaptive or neutral evolution.

### Genome assembly improvement and gene annotation

To identify the genomic mechanism that led to the evolution of differences in lifespan between natural populations of the turquoise killifish (*Nothobranchius furzeri*), we combined the currently available reference genomes^15,18^ into an improved reference turquoise killifish genome assembly. Due to the high repeat content, assembly from short reads required a highly integrated and multi-platform approach. We ran Allpaths-LG with all the available pair-end sequences, producing a combined assembly with a contig N50 of 7.8kb, corresponding to a ∼2kb improvement from the previous versions. Two newly obtained 10X Genomics linked read libraries were used to correct and link scaffolds, resulting in a scaffold N50 of 1.5Mb, i.e. a three-fold improvement from the best previous assembly. With the improved continuity, we assigned 92.2% of assembled bases to the 19 linkage groups using two RAD-tag maps^15^. Gene content assessment using the BUSCO method improved “complete” BUSCOs from 91.43%^15^ and 94.59%^18^ to 95.20%. We mapped Genbank *N. furzeri* RefSeq RNA to the new assembly to predict gene models. The predicted gene model set is 96.1% for “complete” BUSCOs. The overall size of repeated regions (masked regions) is 1.003 Gb, accounting for 66% of the entire genome, i.e. 20% higher than a previous estimate^19^.

### Population genetics of natural turquoise killifish populations

Natural populations of turquoise killifish occur along an aridity gradient in Zimbabwe and Mozambique and populations from more arid regions are associated with shorter captive lifespan^8,14^. A QTL study performed between short-lived and long-lived turquoise killifish populations showed a complex genetic architecture of lifespan (measured as age at death), with several genome-wide loci associated with lifespan differences among long-lived and short-lived populations^15^. To further investigate the evolutionary forces shaping genetic differentiation in the loci associated with lifespan among wild turquoise killifish populations, we performed pooled whole-genome-sequencing (WGS) of killifish collected from four sampling sites within the natural turquoise killifish species distribution, which vary in altitude, annual precipitation and aridity (**Figure S1, Table S1**). Population GNP is located within the Gonarezhou National Park at high altitude and in an arid climate (Koeppen-Geiger classification “BWh”, **Figure S1**), in a region at the outer edge of the turquoise killifish distribution (**Figure S1**)^20-22^, which corresponds to the place of origin of the “GRZ” laboratory strain, which has the shortest lifespan of all laboratory strains of turquoise killifish^14,15^. Population NF414 (MZCS 414) is located in an arid area in the center of the Chefu river drainage in Mozambique (“BWh”, **Figure S1**)^20-22^, and population NF303 (MZCS 303) is located in a semi-arid area in transition to more humid climate zones in the center of the Limpopo river drainage system (Koeppen-Geiger classification “BSh”, **Figure S1**)^20-22^. Altitude among localities ranges from 344 m (GNP) to 68 m (NF303, **Figure S1a** and **Table S1**). The temporary habitat of turquoise killifish populations differs in terms of altitude and aridity, as the ephemeral pools at higher altitude are drained earlier and persist for shorter time, while water bodies in habitats at lower altitude last longer^14^. Population GNP is therefore named “dry”, population NF414 is named “intermediate” and population NF303 “wet” throughout the manuscript.

### High genetic differentiation and contrasting population demography between dry and wet populations

We asked whether populations from dry, intermediate and wet areas, corresponding to shorter and progressively longer lifespan, differ in genetic variability. We calculated genome-wide estimates of average pairwise difference (π) and genetic diversity (*θ*_Watterson_) based on 50kb-non-overlapping sliding windows using PoPoolation^23^. We found that π and *θ*_Watterson_ decrease from wet to dry population (*θ*_Watterson GNP_: 0.0011, *θ*_Watterson NF414_: 0.0036, *θ*_Watterson NF303_: 0.0072; π_GNP_: 0.0009, π_NF414_: 0.0031, and π_NF303_: 0.0054). To infer the genetic distance between the populations, we computed the genome-wide pairwise genetic differentiation between populations using F_ST_^24^. Overall, the genetic differentiation between populations ranged between 0.14 and 0.26 and was the highest between population GNP (dry) and population NF303 (wet) (**Figure 1a**).

**Figure 1.**
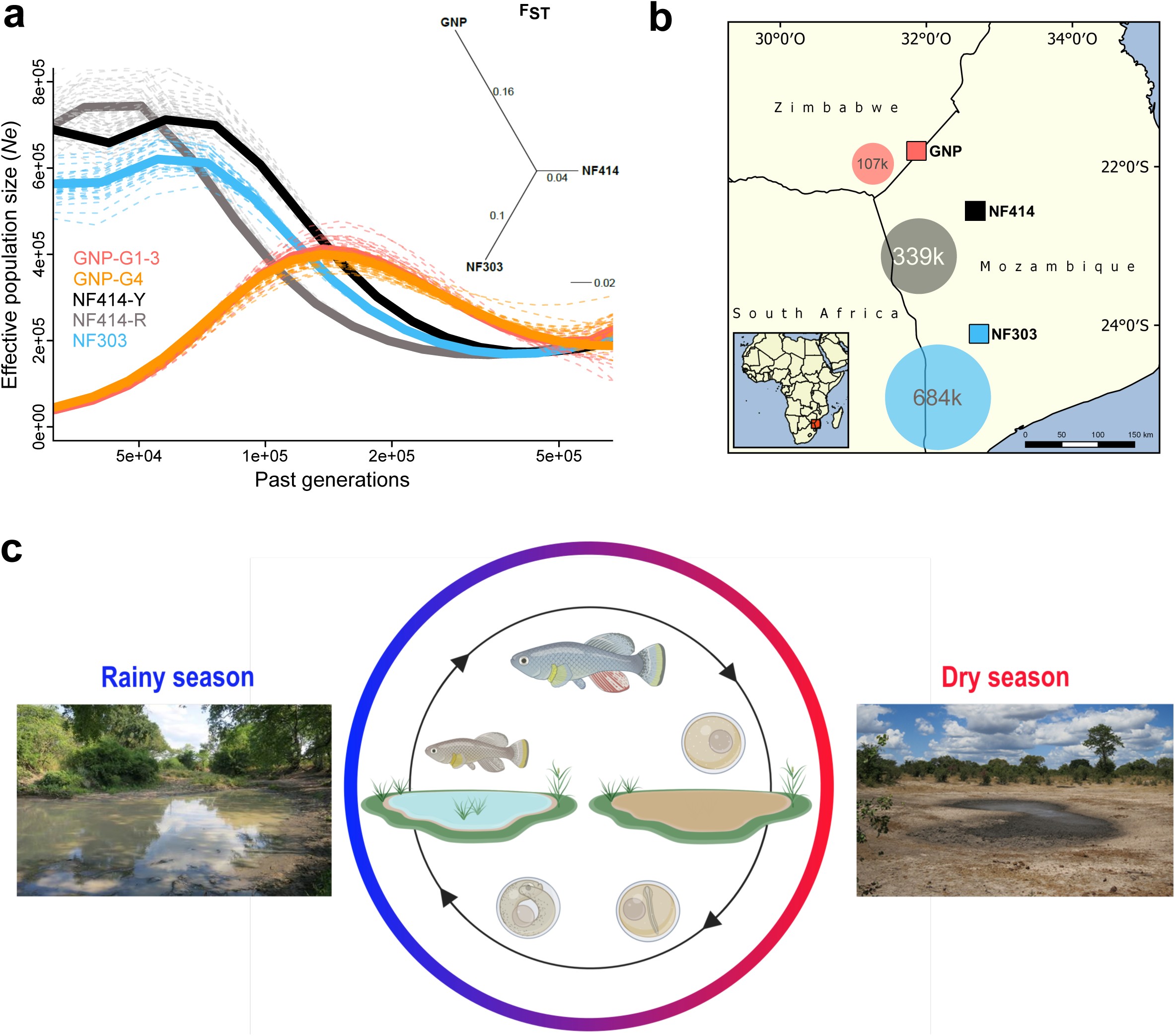
Demography and natural occurence of turquoise killifish populations. a) Inferred ancestral effective population size (*N*_*e*_) (using PSMC’) on y-axis and past generations on x-axis in GNP (red, orange), NF414 (black, grey) and NF303 (blue). Inset: unrooted neighbour joining tree based on pairwise genetic differentiation (F_ST_) values. b) Geographical locations of sampled natural population of turquoise killifish (*Nothobranchius furzeri*). The area of coloured circles represent the estimated effective population size (*N*_*e*_) based on *θ*_Watterson_. c) Natural environment of turquoise killifish and schematic of the annual life cycle. Figure 1 partly made with Biorender®.

Next, we inferred the demographic history of the populations using pairwise sequentially Markovian coalescent (PSMC) by resequencing at high-coverage single individuals for each population^25^. The population GNP (dry) experienced a strong population decline starting approximately 150k generations ago, a result consistent in both sequenced individuals from the two sampling sites (GNP-G1-3 and GNP-G4, **Figure 1a**). In contrast to the demographic history in GNP, we found indications for recent population expansions in populations from the center of the Chefu and Limpopo basins clades. Analysis of population NF414 (intermediate) (**Figure 1a**, NF414-Y and NF414-R) and NF303 (wet) (**Figure 1a**, blue line) shows population expansion until recent time (∼50k generations ago). To infer the effective population size (*N*_*e*_) of the populations, we used the published mutational rate of 2.6321e−9 per base pair per generation for *Nothobranchius* computed via dated phylogeny and *θ*_Watterson_ ^6^. In line with the decrease in genetic diversity from wet to dry population, we found a decrease in *N*_*e*_ estimates (107221.8, 338849.48 and 683693.25 for GNP, NF414 and NF303, respectively; **Figure 1b**). Hence, our findings show that dry populations from the outer edge of the species distribution show lower genetic diversity and smaller effective population size compared to population from intermediate and more wet regions.

### Genetic differentiation among turquoise killifish populations

To test whether regions underlying longevity QTL in turquoise killifish^15,17^ display a genetic signature for positive or purifying selection, we took advantage of the improved turquoise killifish genome assembly and the newly sequenced wild turquoise killifish populations (**Figure 2**). The strongest QTL for lifespan differences among long-lived and short-lived populations mapped on the sex chromosome^15,17^, in proximity to the sex determining locus^15^. To identify a genomic signature of strong selection, we performed an outlier approach based on the pairwise genetic differentiation index (F_ST_). To find highly differentiated regions that may underlie positive selection in natural turquoise killifish populations, we scanned for regions with elevated genetic differentiation between pairs of populations, i.e. exceeding the 0.995 quantile of Z-transformed non-overlapping 50kb sliding windows of F_ST_. To find regions under purifying selection, we scanned for regions with lowered genetic differentiation among populations, i.e. below the 0.005 quantile of Z-transformed non-overlapping 50kb sliding windows of F_ST_ (**TableS7**). The outlier approach did not reveal clear signatures of positive or purifying selection based on genetic differentiation in the four main clusters associated with lifespan in experimental strains of turquoise killifish (**Figure 2**). We then analysed genomic regions carrying signatures of positive and purifying selection in the natural turquoise killifish populations irrespective of the QTL regions (**Figure 2**). The F_ST_ outlier approach led to the identification of several potential regions under divergent selection between populations, in particular between the intermediate and wet populations (**Table S4**) and only two between the dry and wet populations (**Table S5)**. Genes significantly different and within regions of larger genetic differentiation based on Z-transformed non-overlapping sliding windows of F_ST_ were located on chromosomes 6 and 10. The region on chromosome 6 includes the gene *slc8a1*, which contains mutations with significant difference in allele frequencies between the wet and intermediate population (Fisher’s exact test implemented in PoPoolation; adjusted p value < 0.001). The region on chromosome 10 contains four genes: XM_015941868, XM_015941869, *lss* and *hibch.* All genes under the major F_ST_ peak on chromosome 10 showed significant difference in allele frequencies between the intermediate and wet population (Fisher’s exact test; adjusted p value < 0.001) and additionally, *hibch* had significantly different allele frequencies between the dry and wet population (Fisher’s exact test; adjusted p value < 0.001). Genes under F_ST_ peaks between populations that differ in lifespan, are not necessarily causally involved in lifespan differences between populations, as sequence differences could segregate in populations due to population structure and drift. However, to test whether the genes located in genomic regions that are significantly divergent between populations could be functionally involved in age-related phenotypes, we investigated whether gene expression in these genes varied as a function of age. Analysing available turquoise killifish longitudinal RNA-datasets generated in liver, brain and skin^26^, we found that *hibch, lss* and *slc8a1* are differentially expressed between adult and old killifish (**Table S10**, adjusted p value < 0.01). *hibch, lss and slc8a1* are involved in amino acid metabolism^27^, biosynthesis of cholesterol^28^, and proton-mediated accelerated aging^29^, respectively. Gene XM_015956265 (ZBTB14) is the only gene that is an F_ST_ outlier and that is differentially expressed in adult vs. old individuals between at least two populations in all tissues (liver, brain and skin). XM_015956265 encodes a transcriptional modulator with ubiquitous functions, ranging from activation of dopamine transporter to repression of MYC, FMR1 and thymidine kinase promoters^30^. However, although genomic regions that have sequence divergence between turquoise killifish populations contain genes that are differentially expressed during ageing in different tissues, whether any of these genes are causally involved in modulating ageing-related changes between turquoise killifish wild populations still remains to be assessed. Based on the outlier approach, we found two genomic regions with low genetic differentiation between all pairs of populations, indicating strong purifying selection. The first region is located on the sex chromosome and contains the putative sex determining gene *gdf6*^*18*^, which is hence conserved among these populations. This same region also contains *sybu*, a maternal-effect gene associated with the establishment of embryo polarity^31^. The second region under low genetic differentiation is located on chromosome 9 and harbours the genes XM_015965812 (*abi2-like*), *cnot11* and *lcp1*, which are involved in phagocytosis^32^, mRNA degradation^33^ and cell motility^34^, respectively.

**Figure 2.**
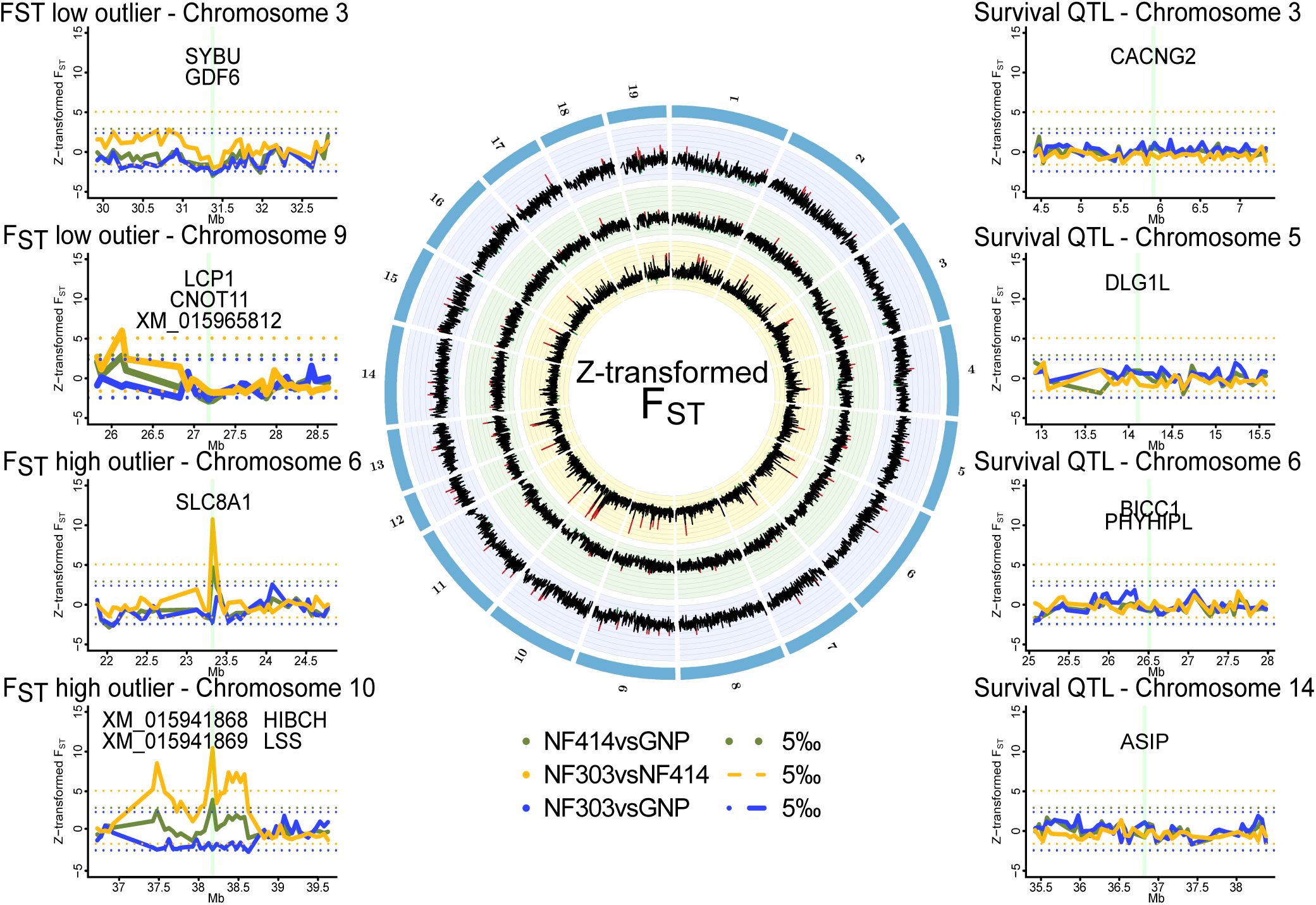
Genomic regions of high and low genetic divergence between pairs of turquoise killifish populations. Left) Genomic regions with high or low genetic differentiation between turquoise killifish populations identified with an F_ST_ outlier approach. Z-transformed F_ST_ values of all pairwise comparison in solid lines, with “NF303vsNF414” in yellow, “NF303vsGNP” in blue, and “NF414vsGNP” in green. The significance thresholds of upper and lower 5‰ are shown in spotted lines with same colour coding. Center) Circos plot of Z-transformed F_ST_ values between all pairwise comparisons with “NF303vsNF414” in the inner circle (yellow), “NF414vsGNP” in the middle circle (green), and “NF303vsGNP” in the outer circle (blue). Right) Pairwise genetic differentiation based on FST in the four main clusters associated with lifespan (QTL from Valenzano et al.^13^).

### Evolutionary origin of the sex chromosome

Since we found reduced genetic differentiation among populations in the chromosomal region containing the putative sex-determining gene in the sex chromosome, we used synteny analysis and the new genome assembly to investigate the genomic events that led to evolution of this chromosomal region (**Figure 3**). We found that the structure of the turquoise killifish sex chromosome is compatible with a chromosomal translocation within an ancestral chromosome and a fusion event between two chromosomes. The translocation event within an ancestral chromosome corresponding to medaka’s chromosome 16 and platyfish’s linkage group 3 led to a repositioning of a chromosomal region containing the putative sex-determining gene *gdf6* (**Figure 3b**). The fusion of the translocated chromosome with a chromosome corresponding to medaka chromosome 8 and platyfish linkage group 16, possibly led to the origin of turquoise killifish sex chromosome. We could hence reconstruct a model for the origin of the turquoise killifish sex chromosome (**Figure 3c**), which parsimoniously places a translocation event before a fusion event. The occurrence of two major chromosomal rearrangements, namely a translocation and a fusion, could have then contributed to suppressing recombination around the sex-determining region^15,35^.

**Figure 3.**
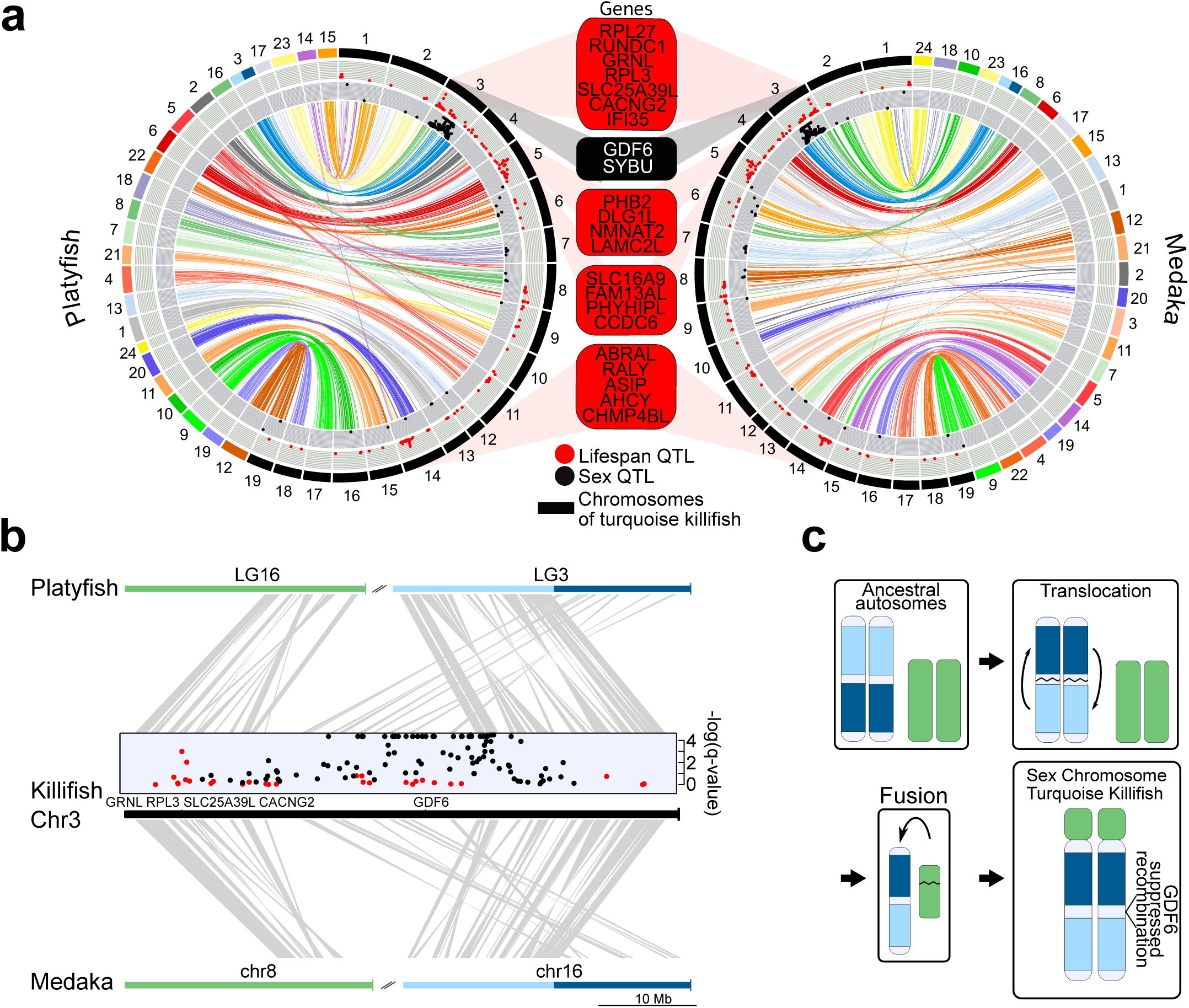
Synteny and sex chromosome evolution in turquoise killifish. a) Synteny circos plots based on 1-to-1 orthologous gene location between the new turquoise killifish assembly (black chromosomes) and platyfish (*Xiphophorus maculatus*, coloured chromosomes, left circos plot) and between the new turquoise killifish assembly (black chromosomes) and medaka (*Oryzias latipes*, coloured chromosomes, right circos plot). Orthologous genes in concordant order are visualized as one syntenic block. Synteny regions are connected via colour-coded ribbons, based on their chromosomal location in platyfish or medaka. If the direction of the syntenic sequence is inverted compared to the compared species, the ribbon is twisted. Outer data plot shows –log(q-value) of survival quantitative trait loci (QTL, ordinate value between 0 and 3.5, every value above 3.5 is visualized at 3.5^13^) and the inner data plot shows –log(q-value) of the sex QTL (ordinate value between 0 and 3.5, every value above 3.5 is visualized at 3.5). Boxes between the two circos plots show genes within the peak regions of the four highest – log(q-value) of survival QTL on independent chromosomes (red box) and the highest association to sex (black box). b) High resolution synteny map between the sex-chromosome of the turquoise killifish (Chr3) with platyfish chromosome 16 and 3 in the upper plot, and between the turquoise killifish and medaka chromosome 8 and 16 (lower plot). The middle plot shows the QTLs for survival and sex along the turquoise killifish sex chromosome. c) Model of sex chromosome evolution in the turquoise killifish. A translocation event within one ancestral autosome led to the emergence of a chromosomal region harbouring a new sex-determining-gene (SDG). The fusion of a second autosome led to the formation of the current structure of the turquoise killifish sex chromosome.

### Relaxed selection in turquoise killifish populations

Since we could not identify specific signatures of genetic differentiation in the genomic regions associated to longevity from previous QTL mapping, we asked whether other evolutionary forces than directional selection may underlie differences in survival among wild turquoise killifish populations. The difference in the recent and past demography between populations (**Figure 1**) led us to ask whether demography could have led to evolutionary changes on genome-wide scale between natural populations. For each population, we calculated the fraction of substitutions driven to fixation by positive selection since divergence from the outgroup species *Nothobranchius orthonotus* (NOR) using the asymptotic McDonald-Kreitman ***α***^36^. Using NOR as an outgroup, we infer the fraction of positive selection by pooling all coding sites (**Figure 4a**). SNPs were called with the program SNAPE^37^, which specifically deals with pooled sequencing. We only included SNPs with a derived frequency between 0.05-0.95 and performed stringent filtering. The asymptotic McDonald-Kreitman ***α*** ranged from −0.21 to −0.01 in comparison to the very closely related sister species *N. orthonotus*, confirming limited genome-wide positive selection since divergence from *N. orthonotus* (**Figure 4a**). The population GNP, located in an arid region at higher altitude and associated with the shortest recorded lifespan, shows the lowest asymptotic McDonald-Kreitman ***α***, as well as lower McDonald-Kreitman ***α*** values throughout all derived frequency bins, potentially suggesting a higher load of slightly deleterious mutations segregating in this population (**Figure 4a**). Using as an outgroup species another annual killifish species, *Nothobranchius rachovii* (NRC), we confirmed the lowest asymptotic McDonald-Kreitman ***α*** value in the dry population GNP (**Figure 4b**). Additionally, using *Nothobranchius rachovii* (NRC) as outgroup species, the asymptotic McDonald-Kreitman ***α*** ranged from −0.06 to 0.23 among populations, indicating that more alleles were driven to fixation by positive selection in the ancestral lineage leading to *Nothobranchius furzeri* and *Nothobranchius orthonotus*. In particular, the wet population NF303 had the highest asymptotic McDonald-Kreitman ***α*** value (**Figure 4b**). Using both *N. orthonotus* and *N. rachovii* as outgroups, we found that the dry GNP population had the lowest McDonald-Kreitman ***α*** values at the low derived frequency bins, potentially consistent with a genome-wide accumulation of slightly deleterious mutations in these isolated populations.

**Figure 4.**
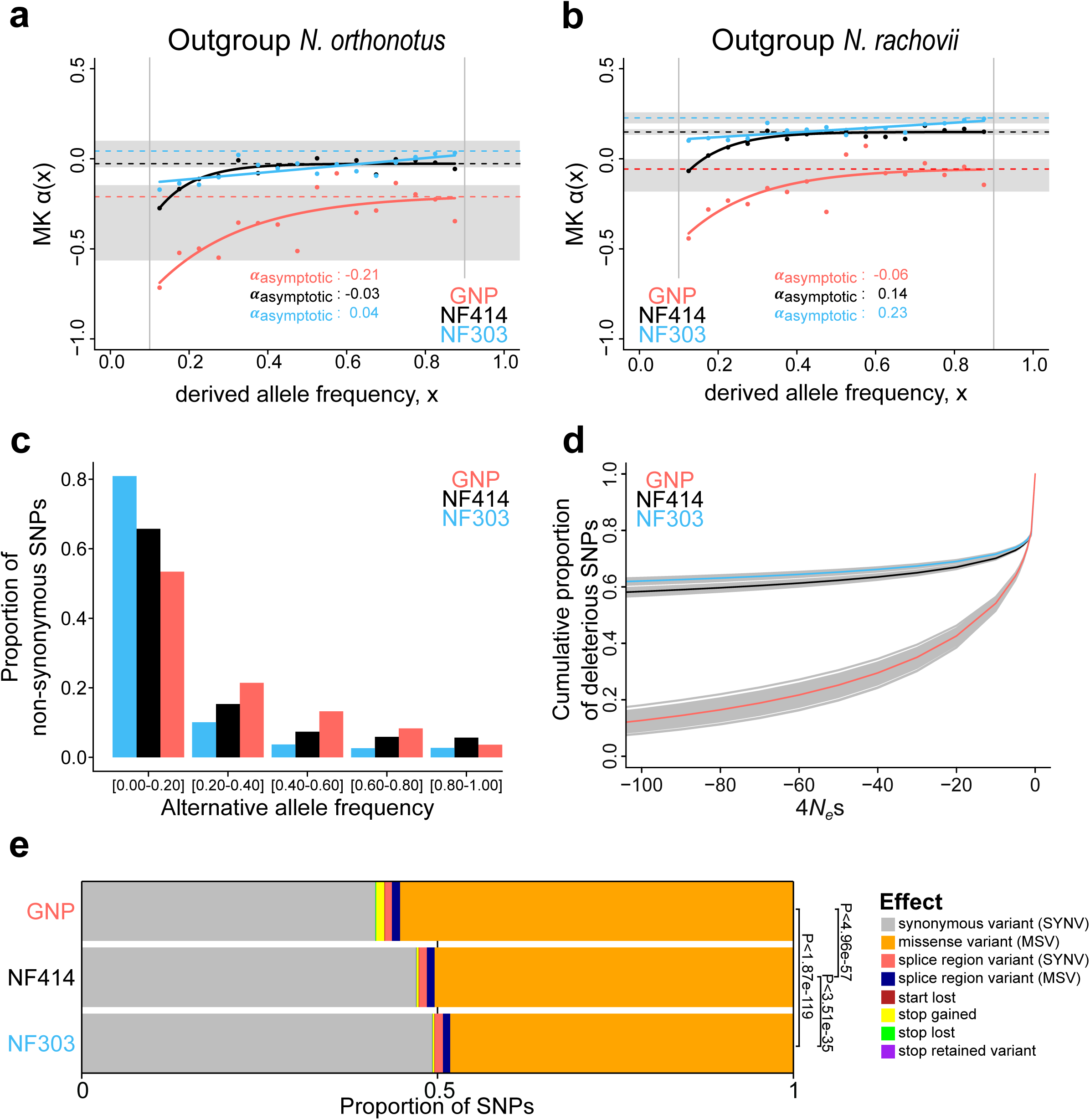
Genome-wide signatures of natural and relaxed selection in turquoise killifish. Asymptotic McDonald-Kreitman alpha (MK ***α***) analysis based on derived frequency bins using as outgroups a) *Nothobranchius orthonotus* and b) *Nothonbranchius rachovii*. Population GNP is shown in red, NF414 in black, and NF303 in blue. c) Proportion of non-synonymous SNPs binned in allele frequencies of non-reference (alternative) alleles for GNP (red), NF414 (black) and NF303 (blue). d) Negative distribution of fitness effects of populations GNP (red), NF414 (black) and NF303 (blue) with cumulative proportion of deleterious SNPs on y-axis and the compound measure of *4N*_*e*_*s* on x-axis. e) Proportion of different effect types of SNPs in coding sequences of all populations. The effect on amino acid sequence for each genetic variant is represented by colours (legend). Significance is based on ratio between synonymous effects to non-synonymous effects (significance based on Chi-square test).

To directly estimate the fitness effect of gene variants associated with each population, we analysed population-specific genetic polymorphisms to assign mutations as beneficial, neutral or detrimental, and determine the distribution of fitness effect (DFE)^38^. Consistently with the overall lower McDonald-Kreitman ***α*** values throughout all derived frequency bins, we found more mutations assigned as the slightly deleterious category in the dry GNP population, compared to the other two populations (indicated by the higher number of deleterious SNPs in proximity to 4N_e_S ∼ 0 in the GNP population, **Figure 4d, TableS8**). To further infer the effect of the putative deleterious mutations on protein function, we used the new turquoise killifish genome assembly as a reference and adopted an approach that, by analysing sequence polymorphism among populations, predicts functional consequences at the protein level^39^. We found that the proportion of mutations causing a change in protein function is significantly larger in the GNP population compared to populations NF414 and NF303 (Chi-square test: P_GNP-NF303_<1.87e-119, P_GNP-NF414_< 4.96e-57, P_NF303-NF414_< 3.51e-35, **Figure 4e**). Additionally, the mutations with predicted deleterious effects on protein function reached also higher frequencies in the dry population GNP (**Figure 4c**). To further investigate the impact of mutations on protein function, we calculated the Consurf ^40-43^ score, which determines the evolutionary constraint on an amino acid, based on sequence conservation. Mutations at amino acid positions with high Consurf score (i.e. otherwise highly conserved) are considered to be more deleterious. We found that the dry population GNP had a significantly higher mean Consurf score for mutations at non-synonymous sites in frequency bins from 5%-20% up to 40%-60%, compared to populations NF414 (intermediate) and NF303 (wet) (**Figure S2**). The mutations in the dry GNP population had significantly higher Consurf scores than the other populations using both outgroup species *N. orthonotus* and *N. rachovii* (**Figure S2**). Upon exclusion of potential mutations at neighbouring sites (CMD: codons with multiple differences), CpG hypermutation and genes containing mutations with highly detrimental effect on protein function based on SnpEFF analysis, the dry population GNP had higher mean Consurf score at the low frequency bin (**Figure S2, Table S11-12**). To note, we also found a significantly higher average Consurf score at synonymous sites in GNP at low derived frequencies (Figure S2, Supplementary Table S11 and S12), possibly suggestive of an overall higher mutational rate in GNP.

### Relaxation of selection in age-related disease pathways

Computing the gene-wise direction of selection (DoS)^44^ index, which enables to score the strength of selection based on the count of mutations in non-synonymous and synonymous sites, we found support to the hypothesis that the dry, short-lived population GNP has significantly more slightly deleterious mutations segregating in the population, compared to the populations NF414 and NF303 (**Figure 5a**, Median NOR: GNP: −0.17, NF414: −0.02, NF303: −0.01; Median NRC: GNP: −0.14, NF414: 0.00, NF303: 0.00; Wilcoxon rank sum test: NOR: P_GNPNF303_<2.21e-105, P_GNP-NF414_< 1.19e-76, P_NF303-NF414_< 1.39e-06; NRC: P_GNP-NF303_<4.61e-179, P_GNP-NF414_< 1.42e-100, P_NF303-NF414_< 5.96e-22), indicating that purifying selection is relaxed in GNP. We calculated DoS in all populations using independently as outgroup species *N. orthonotus* and *N. rachovii* (**Figure 5a**).

**Figure 5.**
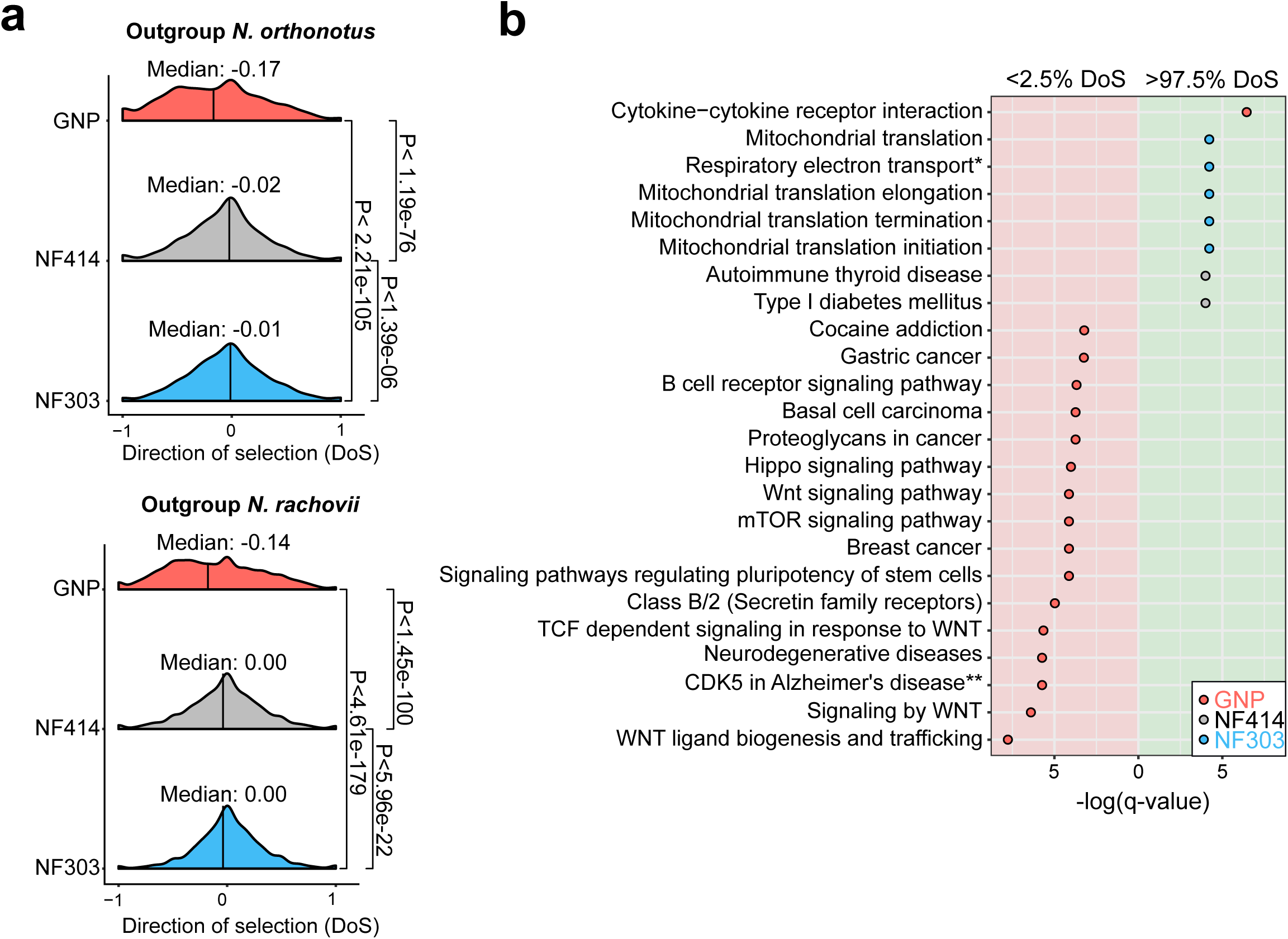
Pathway enrichment in genes under adaptive and neutral evolution in turquoise killifish populations. a) Distribution of direction of selection (DoS) represented with median of distribution for population GNP (red), NF414 (grey) and NF303 (blue). Left panel shows DoS distribution computed using *Nothobranchius orthonotus* as outgroup and right panel shows DoS distribution computed using *Nothobranchius rachovii* as outgroup. Significance based on Wilcoxon-Rank-Sum test. b) Pathway over-representation analysis of genes below the 2.5% level of gene-wise DoS values are shown with red background and above the 97.5% level of gene-wise DoS values are shown with green background. Only pathway terms with significance level of FDR corrected q-value < 0.05 are shown (in -log(q-value)). Terms enriched in population GNP have red dots, enriched in population NF414 have black dots, and enriched in population NF303 have blue dots, respectively. *ATP synthesis by chemiosmotic coupling, and heat production by uncoupling proteins. ** Deregulated CDK5 triggers multiple neurodegenerative pathways in Alzheimer’s disease models.

To assess whether specific biological pathways were significantly more impacted by the accumulation of slightly deleterious mutations, we performed pathway overrepresentation analysis. We found a significant overrepresentation in the lower 2.5^th^ DoS (i.e. genes under relaxation of selection) in the GNP population for pathways associated with age-related diseases, including *gastric cancer, breast cancer, neurodegenerative disease, mTOR signalling* and WNT signalling (q-value <0.05, **Figure5b, Table S9**). Overall, relaxed selection in the dry GNP population affected accumulation of deleterious mutations in age-related and in the WNT pathway. Analysing the pathways affected by genes within the upper 2.5^th^ DoS values – corresponding to genes undergoing adaptive evolution – we found a significant enrichment for mitochondrial pathways – potentially compensatory^6^ – in population NF303 (**Figure5b, Table S9**). Overall, our results show that differences in effective population size among wild turquoise killifish are associated with an extensive relaxation of purifying selection, significantly affecting genes involved in age-related diseases, and which could have cumulatively contributed to reducing survival.

## Discussion

The turquoise killifish (*Nothobranchius furzeri*) is the shortest-lived known vertebrate and while its natural populations show similar timing for sexual maturation, exhibit differences in lifespan along a cline of altitude and aridity in south-eastern Africa^8,14^. Here we generate an improved genome assembly (NFZ v2.0) in turquoise killifish (*Nothobranchius furzeri*) and study the evolutionary forces shaping genome evolution among natural populations.

Using the new turquoise killifish genome assembly and synteny analysis with medaka and platyfish, we reconstructed the origin of the turquoise killifish sex chromosome, which appears to have evolved through two independent chromosomal events, i.e. a fusion and a translocation event.

Using the new genome assembly and pooled sequencing of natural turquoise killifish populations, we found that genetic differentiation among populations of the short-lived turquoise killifish is consistent with differences in demographic constraints. While we found that strong purifying selection maintains low genetic diversity among populations at genomic regions underlying key species-specific traits, such as in proximity to the sex-determining region, demography and genetic drift largely shape genome evolution, leading to relaxation of selection and the accumulation of deleterious mutations. We showed that isolated populations from an arid region, dwelling at higher altitude and characterised by shorter lifespan, experienced extensive population bottlenecking and a sharp decline in effective population size. We found that relaxation of selection in highly drifted populations significantly affected the accumulation of deleterious gene variants in pathways associated with neurodegenerative diseases and WNT-signalling (**Figure 5**). While simple traits, such as male tail colour and sex have a simple genetic architecture among turquoise killifish populations^15,35^, we find that the complex genetic architecture of lifespan differences among killifish populations^15^ is entirely compatible with genome-wide relaxation of selection. Additionally, the absence of genomic signature of positive selection in genomic regions underlying survival QTL in killifish suggest that, rather than directional selection, the neutral accumulation of deleterious mutations in short-lived populations may be the evolutionary mechanism underlying survival differences among turquoise killifish populations. The “antagonistic pleiotropy” evolutionary theory of ageing states that positive selection could lead to the fixation of gene variants that, while overall beneficial for fitness, could reduce survival and reproductive capacity in late life^45^. The lack of genomic signature of positive selection at the genomic regions underlying survival QTL in turquoise killifish rather suggests that accumulation of deleterious mutations may have played a key role in shaping genome and phenotype differences among natural turquoise killifish populations. Historical fluctuations in the size of natural turquoise killifish populations, especially in isolated and populations living in more arid and elevated habitats, cause decreased efficiency of the strength of natural selection, ultimately contributing to increased load of deleterious gene variants, preferentially in genes associated with ageing-related diseases and in the WNT pathway.

Our findings highlight the role of demographic constraints in shaping life history within species.

## Materials and Methods

### Merging and Improvement of the Turquoise killifish genome assembly

#### 10x Genomics read clouds

A single GRZ male individual was sacrificed with MS222 (Sigma-Aldrich, Steinheim, Germany). Blood was drawn from the heart and high molecular weight DNA was isolated with Qiagen MagAttract kit following manufacturer’s instructions. Gemcode v2 DNA library generation was performed by Novogene (Beijing, China). Briefly, a proportion of the sample was run on a pulse field agarose gel to confirm high molecularity > 100kb. Based on a genome size estimate of 1.54Gb (half of human genome), 0.6ng of DNA was used to construct 2 Gemcode libraries, sequenced on two HiSeq X lanes to obtain a raw coverage of approximately 60X each. The reported input molecular length by SuperNova^46^ was 118kb for library 1 and 60.73kb for library 2. Both libraries were used to correct and scaffold the Allpath-LG assembly (see below), and library 1 was also de novo assembled with the SuperNova assembler v.2 with default parameters. The SuperNova assembly totaled 802.6Mb, with a contig N50 of 19.65kb, scaffolded into 6.78 thousand scaffolds with an N50 of 3.83Mb. Despite high continuity, however, the BUSCO^47^ metrics are much lower than the Allpath-LG assemblies.

#### Nanopore long reads

DNA was extracted from a single GRZ male individual’s muscle tissue by griding in liquid nitrogen followed by phenol-chlorofom extraction (Sigma). The rapid sequencing kit (SQK-RAD004) and the ligation kit (SQK-LSK108) were sued to prepare 6 libraries and were sequenced on 6 MinION flow cells (R9.4.1). These runs yielded a total of 3.3 Gb of sequences after trimming and correction by HALC^48^. For correction, Allpath-LG contigs (see below) and short reads from the 10X genomic run were used.

#### Allpath-LG assembly

Two independent short read datasets were previous collected for the GRZ strain of *Nothobranchius furzeri*. Allpath-LG^49^ was used on the pooled datasets. Together, 4 illumina short read pair-end libraries with a fragment size distribution from 158bp to 179bp were used to construct the contigs (sequence coverage 191.9X, physical coverage 153.5X), and 22 pair-end and mate pair libraries distributed at 92bp, 135bp, 141bp, 176bp, 267bp, 2kb, 3kb 5kb and 10kb were used for the scaffolding step (sequence coverage 135.7X, physical coverage 453.8X). The published BAC library ends^18^ with an insert size of 112kb were also included in the ALLPaths-LG run (physical coverage 0.6X). The resulting assembly has a total contig length of 823,583,106bp distributed in 151,307 contigs > 1kb, with an N50 of 7.8kb. The total scaffold length is 943,793,727bp distributed in 7830 scaffolds with an N50 of 421kb (with gaps). The resulting assembly was further scaffolded by ARCS v1.0^50^ + LINKS v1.8.5^51^ with the following parameters: arcs -e 50000 -c 3 -r 0.05 -s 98 and LINKS -m -d 4000 -k 20 -e 0.1 -l 3 -a 0.3 -t 2 -o 0 -z 500 -r -p 0.001 -x 0. This increased the scaffold N50 to 1.527 Mb. Next, scaffolds were assigned to the RAD-tag linkage map^15^ collected from a previous study with Allmaps ^52^, using equal weight for the two independent mapping crosses. This procedure assigned 90.6% of the assembled bases in 1131 scaffolds to 19 linkage groups, in which 76.6% can be oriented. Misassemblies were corrected with the 10X genomic read cloud. Read clouds were mapped to the preliminary assembly with longranger v2.1.6 using default parameters, and a custom script was used to scan for sudden drops in barcode shares along the assembled linkage groups. The scaffolds were broken at the nearest gap of the drop in 10x barcodes. The same ARCS + LINKS pipeline was again run on the broken scaffolds, increasing the scaffold N50 to 1.823 Mb. Next, BESST_RNA (https://github.com/ksahlin/BESST_RNA) was used to further scaffold the assembly with RNASeq libraries, Allmaps was again used to assign the fixed scaffolds back to linkage groups, increasing the assignable bases to 92.2% (879Mb) with 80.3% (765Mb) with determined orientation. The assembly was again broken with longranger and reassigned to LG with Allmaps, and the scaffolds were further partitioned to linkage groups due to linkage of some left-over scaffolds with an assigned scaffold. Each partitioned scaffold groups were subjected to the ARCS + LINKS pipeline again, to constraint the previously unassigned scaffolds onto the same linkage group. Allmaps was run again on the improved scaffolds, resulting in 94.5% (903.4Mb) of bases assigned and 89.1% (852Mb) of bases oriented. Longranger was run again, visually checked and compared with the RADtag markers. Eleven mis-oriented positions were identified and corrected. Gaps were further patched by GMCloser^53^ with ∼2X of nanopore long reads corrected by HALC using BGI500 short PE reads with the following parameters: gmcloser --blast --long_read --lr_cov 2 -l 100 -i 466 -d 13 --min_subcon 1 --min_gap_size 10 --iterate 2 -- mq 1 -c. The corrected long reads not mapped by GMCloser were assembled by CANU ^54^ into 7.9Mb of sequences, which are likely unassigned repeats.

#### Meta Assembly

Five assemblies were integrated by MetAssembler^55^ in the following order (ranked by BUSCO scores) using a 20kb mate pair library: 1) The improved Allpaths-LG assembly assigned to linkage groups produced in this study, 2) A previously published assembly with Allpaths-LG and optical map^18^ 3) A previously published assembly using SGA^15^, 4) The SuperNova assembly with only 10x Genomic reads and 5) Unassigned nanopore contigs from CANU. The final assembly NFZ v2.0 has 911.5Mb of scaffolds assigned to linkage groups. Unassigned scaffolds summed up to 142.2Mb, yielding a total assembly length of 1053.7Mb, approximately 2/3 of the total genome size of 1.53Gb. The final assembly has 95.2% complete and 2.24% missing BUSCOs.

#### Mapping of NCBI genbank gene annotations

RefSeq mRNAs for the GRZ strain (PRJNA314891, PRJEB5837) were downloaded from GenBank^56^, and aligned to the assembly with Exonerate^57^. The RefSeq mRNAs have a BUSCO score of 98.0% complete, 0.9% missing. The mapped gene models resulted in a BUSCO score of 96.1% complete, 2.1% missing.

#### Pseudogenome assembly generation

The pseudogenomes for *Nothobranchius orthonotus* and *Nothobranchius rachovii* were generated from sequencing data and the same method used in Cui *et al*. ^6^. Briefly, the sequencing data were mapped to the NFZ v2.0 reference genome by BWA-mem v0.7.12 in PE mode^58,59^. PCR duplicates were marked with MarkDuplicates tool in the Picard (version 1.119, http://broadinstitute.github.io/picard/) package. Reads were realigned around INDELs with the IndelRealigner tool in GATK v3.4.46^60^. Variants were called with SAMTOOLS v1.2^61^ mpileup command, requiring a minimal mapping quality of 20 and a minimal base quality of 25. A pseudogenome assembly was generated by substituting reference bases with the alternative base in the reads. Uncovered regions, INDELs and sites with >2 alleles were masked as unknown “N”. The allele with more supporting reads was chosen at biallelic sites.

#### Mapping of longevity and sex quantitative trait loci

The quantitative trait loci (QTL) markers published in Valenzano *et al.*^15^ were directly provided by Dario Riccardo Valenzano. In order to map the markers associated to longevity and sex, a reference database was created using BLAST^62^. The nucleotide database was created with the new reference genome of *N. furzeri* (NFZ v2.0). Subsequently, the QTL marker sequences were mapped to the database. Only markers with full support for the total length of 95 bp were considered as QTL markers.

#### Synteny analysis

Synteny analysis was performed using orthologous information from Cui *et al*. ^6^ determined by the UPhO pipeline^63^. For this, the 1-to-1 orthologous gene positions of the new turquoise killifish reference genome (NFZ v2.0) were compared to two closely related teleost species, *Xiphophorus maculatus* and *Oryzias latipes*. Result were visualized using Circos^64^ for the genome-wide comparison and the *genoPlotR* package^65^ in R for the sex chromosome synteny analysis. Synteny plots for orthologous chromosomes of *Xiphophorus maculatus* and *Oryzias latipes* were generated with Synteny DB (http://syntenydb.uoregon.edu) ^66^.

#### Koeppen-Geiger index and bioclimatic variables

The Koeppen-Geiger classification data was taken from Peel *et al.*^67^ and the altitude, precipitation per month, and the bioclimatic variables were obtained from the Worldclim database (v2.0^68^). The monthly evapotranspiration was obtained from Trabucco and Zomer^69^. Aridity index was calculated based on the sum of monthly precipitation divided by sum of monthly evapotranspiration. Maps in FigureS1 were generated with QGIS version 2.18.20 combined with GRASS version 7.4^70^, the Koeppen-Geiger raster file, data from Natural Earth, and the river systems database from Lehner *et al.*^71^.

#### DNA isolation and pooled population sequencing

The ethanol preserved fin tissue was washed with 1X PBS before extraction. Fin tissue was digested with 10 µg/mL Proteinase K (Thermo Fisher) in 10 mM TRIS pH 8; 10mM EDTA; 0.5 SDS at 50°C overnight. DNA was extracted with phenol-chloroform-isoamylalcohol (Sigma) followed by a washing step with chloroform (Sigma). Next, DNA was precipitated by adding 2.5 volume of chilled 100% ethanol and 0.26 volume of 7.5M Ammonium Acetate (Sigma) at −20°C overnight. DNA was collected via centrifugation at 4°C at 12000rpm for 20 minutes. After a final washing step with 70% ice-cold ethanol and air drying, DNA was eluted in 30µl of nuclease-free water. DNA quality was checked on 1 agarose gels stained with RotiSafe (Roth) and a UV-VIS spectrometer (Nanodrop2000c, Thermo Scientific). DNA concentration was measured with Qubit fluorometer (BR dsDNA Assay Kit, Invitrogen). For each population, the DNA of the individuals were pooled at equimolar contribution (GNP_G1_3, GNP_G4 N=29; NF414, NF303 N=30). DNA pools were given to the Cologne Center of Genomic (CCG, Cologne, Germany) for library preparation. The total amount of DNA provided to the sequencing facility was 3.2 µg per pooled population sample. Libraries were sequenced with 150bp × 2 paired-ends on the HiSeq4000. Sequencing of pooled samples resulted in a range of 419 - 517 million paired-end reads for each population (**Table S2**).

#### Mapping of pooled sequencing reads

Raw sequencing reads were trimmed using Trimmomatic-0.32 (ILLUMINACLIP:illumina-adaptors.fa:3:7:7:1:true, LEADING:20, TRAILING:20, SLIDINGWINDOW:4:20, MINLEN:50^72^. Data files were inspected with FastQC (version 0.11.22, https://www.bioinformatics.babraham.ac.uk/projects/fastqc/). Trimmed reads were subsequently mapped to the reference genome with BWA-MEM v0.7.12^58,59^. The SAM output was converted into BAM format, sorted, and indexed via SAMTOOLS v1.3.1^61^. Filtering and realignment was conducted with PICARD v1.119 (http://broadinstitute.github.io/picard/) and and GATK^60^. Briefly, the reads were relabelled, sorted, and indexed with AddOrReplaceReadGroups. Duplicated reads were marked with the PICARD feature MarkDuplicates and reads were realigned with first creating a target list with RealignerTargetCreator, second by IndelRealigner from the GATK suite. Resulting reads were again sorted and indexed with SAMTOOLS. For population genetic bioinformatics analyses the BAM files of the pooled populations were converted into the required MPILEUP format via the SAMTOOLS mpileup command. Low quality reads were excluded by setting a minimum mapping quality of 20 and a minimum base quality of 20. Further, possible insertion and deletions (INDELs) were identified with *identifygenomic-indel-regions.pl* script from the PoPoolation package^23^ and were subsequently removed via the *filter-pileup-by-gtf.pl* script^23^. Coding sequence positions that were identified to be putative ambiguous were removed by providing the *filter-pileup-by-gtf.pl* script a custom modified GTF file with the corresponding coordinates. After adapter and quality filtering, mapping to the newly assembled reference genome resulted in mean genome coverage of 35x, 39x, and 47x for the population NF303, NF414, and GNP, respectively (**Table S2**).

#### Merging sequencing reads of populations from the Gonarezhou National Park

Population GNP consists of two sampling sites (GNP-G1_3, GNP-G4) with very low genetic differentiation (**Figure S1c, Table S3**). Sequencing reads of the two populations from the Gonarezhou National Park (GNP) were combined used the SAMTOOLS ‘merge’ command. The populations GNP-G1-3 and GNP-G4 were merged together and this population was subsequently denoted as GNP.

#### Estimating genetic diversity

Genetic diversity in the populations was estimated by calculating the nucleotide diversity π^73^ and Wattersons’s estimator θ^74^. Calculation of π and θ was done with a sliding window approach by using the *Variance-sliding.pl* script from the PoPoolation program^23^. Non-overlapping windows with a length of 50 kb with a minimum count of two per SNP, minimum quality of 20 and the population specific haploid pool size were used (GNP=116; NF414=60; NF303=60). Low covered regions that fall below half the mean coverage of each population were excluded (GNP=23; NF414=19; NF303=18), as well as regions that exceed a two times higher coverage than the mean coverage (GNP=94; NF414=77; NF303=70). The upper threshold is set to avoid regions with possible wrong assemblies. Mean coverage was estimated on filtered MPILEUP files. Each window had to be at least covered to 30% to be included in the estimation.

#### Estimation of effective population size

Wattersons’s estimator of θ^74^ is referred to as the population mutation rate. The estimate is a compound parameter that is calculated as the product of the effective population size (*N*_*e*_), the ploidy (2p, with p is ploidy) and the mutational rate µ (θ = 2p*N*_*e*_µ). Therefore, *N*_*e*_ can be obtained when θ, the ploidy and the mutational rate µ are known. The turquoise killifish is a diploid organism with a mutational rate of 2.6321e−9 per base pair per generation (assuming one generation per year in killifish ^6^ and θ estimates were obtained with PoPoolation (see Section 2.1.2)^23^.

#### Estimating population differentiation index F_ST_

The filtered and realigned BAM files of each population were merged into a single pileup file with SAMTOOLS mpileup, with a minimum mapping quality and a minimum base quality of 20. The pileup was synchronized using the *mpileup2sync.jar* script from the PoPoolation2 program ^24^. Insertions and deletions were identified and removed with the *identify-indel-regions.pl* and *filter-sync-by-gtf.pl* scripts of PoPoolation2 ^24^. Again, coding sequence positions that were identified to be putative ambiguous were removed by providing the *filter-pileup-by-gtf.pl* script a custom modified GTF file with the corresponding coordinates. Further a synchronized pileup file for genes only were generated by providing a GTF file with genes coordinates to the *create-genewise-sync.pl* from PoPoolation2^24^. F_ST_ was calculated for each pairwise comparison (GNP vs NF303, GNP vs NF414, NF414 vs NF303) in a genome-wide approach using non-overlapping sliding windows of 50kb with a minimum count of four per SNP, a minimum coverage of 20, a maximum coverage of 94 for GNP, 77 for NF414, and 70 for NF303 and the corresponding pool size of each population (N= 116; 60; 60). Each sliding window had to be at least covered to 30% to be included in the estimation. The same thresholds, except the minimum covered fraction, with different sliding window sizes were used to calculate the gene-wise F_ST_ for the complete gene body (window-size of 2000000, step-size of 2000000) and single SNPs within genes (window-size of 1, step-size of 1). The non-informative positions were excluded from the output. Significance of allele differences per base-pair within the gene-coordinates were calculated with the fisher’s exact test implemented in *the fisher-test.pl* script of PoPoolation2 ^24^. Calculation of unrooted neighbor joining tree based on the genome-wide pairwise F_ST_ averages was performed with the ape package in R^75^.

#### Detecting signatures of selection based on F_ST_ outliers

For F_ST_-outlier detection, the pairwise 50kb-window F_ST_-values for each comparison were Z-transformed (Z_FST_). Next, regions potentially under strong selection were identified by applying an outlier approach. Outliers were identified as non-overlapping windows of 50 kb within the 0.5% of lowest and highest genetic differentiation per comparison. To reduce the number of false-positive results, the outlier threshold was chosen at 0.5% highest and lowest percentile of each pairwise genetic differentiation^76,77^. To find candidate genes within windows of highest differentiation, a total of three selection criteria were used. First, the window-based Z_FST_ value had to be above the 99.5^th^ percentile of pairwise genetic differentiation. Second, the gene F_ST_ value had to be above the 99.5^th^ percentile of pairwise genetic differentiation and last, the gene needed to include at least one SNP with significant differentiation based on Fisher’s exact test (calculated with PoPoolation2^24^; P<0.001, Benjamini-Hochberg corrected P-values ^78^).

#### Identifying polymorphic sites

SNP calling was performed with Snape^37^. The program requires information of the prior nucleotide diversity *θ*. Hence, the initial values of nucleotide diversity obtained with PoPoolation were used. Snape was run with folded spectrum and prior type informative. As Snape requires the MPILEUP format, the previously generated MPILEUP files were used. SNP calling was separately performed on coding and non-coding parts of the genome. Therefore, each population MPILEUP file was filtered by coding sequence position with the *filter-pileup-by-gtf.pl* script of PoPoolation. For coding sequences the --keep-mode was set to retain all coding sequences. The non-coding sequences were obtained by using the default option and thus discarding the coding sequences from the MPILEUP file. Snape produces a posterior probability of segregation for each position. The posterior probability of segregation was used to filter low-confidence SNPs and indicated in the specific section.

#### Divergence and polymorphisms in 0-fold and 4-fold sites

Polarization of synonymous sites (four-fold degenerated sites) and non-synonymous sites (zero-fold degenerated sites) was done using the pseudogenomes of outgroups *Nothobranchius orthonotus* and *Nothobranchius rachovii*. For each population the genomic information of the respective pseudogenome was extracted with bedtools getfasta command ^79,80^ and the derived allele frequency of every position was inferred with a custom R script. Briefly, only sites with the bases A, G, T or C in the outgroup pseudogenome were included and checked whether the position has an alternative allele in each of the investigated populations. Positions with an alternative allele present in the population data were treated as possible divergent or polymorphic sites. The derived frequency was determined as frequency of the allele not present in the outgroup. Occasions with an alternate allele present in the population data were treated as possible divergent or polymorphic sites. Divergent sites are positions in the genome were the outgroup allele is different from the allele present in the population. Polymorphic sites are sites in the genome that have more than one allele segregating in the population. Only biallelic polymorphic sites were used in this analysis. The DAF was determined as frequency of the allele not shared with the respective outgroups. In general, positions with only one supporting read for an allele were treated as monomorphic sites. SNPs with a DAF < 5% or > 95% were treated as fixed mutations. Further filtering was done based on the threshold of the posterior probability of >0.9 calculated with Snape (see previous subsection), combined with a minimum and maximum coverage threshold per population (GNP: 24, 94; NF414:19, 77; NF303: 18, 70).

#### Asymptotic McDonald-Kreitman α

The rate of substitutions that were driven to fixation by positive selection was evaluated with an improved method based on the McDonald-Kreitman test^81^. The test assumes that the proportion of non-synonymous mutations that are neutral has the same fixation rate as synonymous mutations. Therefore, under neutrality the ratio between non-synonymous to synonymous substitutions (Dn/Ds) between species is equal to the ratio of non-synonymous to synonymous polymorphisms within species (Pn/Ps). If positive selection takes place, the ratio between non-synonymous to synonymous substitutions between species is larger than the ratio of non-synonymous to synonymous polymorphism within species^81^. The concept behind this is that the selected variant reaches fixation in a shorter time than by random drift. Therefore, the selected variant increases Dn, not Pn. The proportion of non-synonymous substitutions that were fixed by positive selection (α) was estimated with an extension of the McDonald-Kreitman test^82^. Due to the presence of slightly deleterious mutations the estimate of α can be underestimated. For this reason, the method used by Messer and Petrov was implemented to calculate α as a function of the derived allele frequency x^36,83^. With this method the true value of α can be inferred as the asymptote of the function of α. Additionally, the value of α(x) for low derived frequencies should give an estimate of the number of slightly deleterious mutations that segregate in the population.

#### Direction of selection (DoS)

To further investigate the signature of selection, the direction of selection (DoS) index for every gene was calculated^44^. DoS standardizes α to a value between −1 and 1. A positive value of DoS indicates adaptive evolution (positive selection) and a negative value indicates the segregation of slightly deleterious alleles, therefore weaker purifying selection^44^. This ratio is undefined for genes without any information about polymorphic or substituted sites. Therefore, only genes with at least one polymorphic and one substituted site were included.

#### Inference of distribution of fitness effects

The distribution of fitness effects (DFE) was inferred using the program polyDFE2.0^38^. For this analysis the unfolded site frequency spectra (SFS) of non-synonymous (0-fold) and synonymous sites (4-fold) were projected into 10 chromosomes for each population. Information about the fixed derived sites was included in this analysis (using *Nothobranchius orthonotus*). PolyDFE2.0 estimates either the full DFE, containing deleterious, neutral and beneficial mutations, or only the deleterious DFE. The best model for each population was obtained using a model testing approach with three different models implemented in PolyDFE2.0 (Model A, B, C). Due to possible biases from erroneous polarization or unknown demography, runs accounting for polarization errors and demography (+eps, +r) were included. Initial parameters were automatically estimated with the –e option, as recommended. To ensure that the parameter space is explored thoroughly, the basin hopping option was applied with a maximum of 500 iterations (-b). The best model for each population was chosen based on the Akaike Information Criterion (AIC). Confidence intervals were generated by running 200 bootstrap datasets with the same parameters used to infer the best model.

#### Variant annotation

Classification of changes in the coding-sequence (CDS) was done with the variant annotator SnpEFF ^39^. The new genome of *Nothobranchius furzeri* (NFZ v2.0) was implemented to the SnpEFF pipeline. Subsequently, a database for variant annotation with the genome NFZ v2.0 FASTA file and the annotation GTF file was generated. For variant annotation the population specific synonymous and non-synonymous sites with a change in respect to the reference genome NFZ v2.0 were used to infer the impact of these sites. The possible annotation impact classes were low, moderate, and high. SNPs with a frequency below 5% or above 95% were excluded for this analysis. To be consistent with the analysis of the distribution of fitness effects, only positions also found to be present in the *N. orthonotus* pseudogenome were considered. Positions with warnings in the variant annotation were removed.

### Consurf analysis

The Consurf score was calculated accordingly to the method used in Cui *et al.* ^6^. We used the Consurf ^40-43^ package to assign each AA a conservation score based on the evolutionary rate in homologs of other vertebrates. Consurf scores were estimated for 12575 genes of *N. furzeri* and synonymous and non-synonymous genomic positions were matched with the derived allele frequency of *N. orthonotus* and *N. rachovii*, respectively. The derived frequencies were binned in five bins and we used pairwise Wilcoxon rank sum test to assess significance after correcting for multiple testing (Benjamini & Hochberg adjustment) between each subsequent bin per population and matching bins between populations.

#### Over-representation analysis

Gene ontology (GO) and pathway overrepresentation analysis was performed with the online tool ConsensusPathDB (http://cpdb.molgen.mpg.de;version34)^84^ using “KEGG” and “REACTOME” databases. Briefly, each gene present in the outlier list was provided with an ENSEMBL human gene identifier^85^, if available, and entered as the target list into the user interface. All genes included in the analysis and with available human ENSEMBL identifier were used as the background gene list. ConsensusPathDB maps the entries to the databases and calculates the enrichment score for each entity by comparing the proportion of target genes in the entity over the proportion of background genes in the entity. For each of the enrichment a P-value is calculated based on a hyper geometric model and is corrected for multiple testing using the false discovery rate (FDR). Only GO terms and pathways with more than two genes were included. Overrepresentation analysis was performed on genes falling below the 2.5th percentile or above the 97.5th percentile thresholds. The percentiles for either F_ST_ or DoS values were calculated with the quantile() function in R.

### Statistical analysis and data processing

Statistical analyses were performed using R studio version 1.0.136 (R version 3.3.2^86^) on a local computer and R studio version 1.1.456 (R version 3.5.1) in a cluster environment at the Max-Planck-Institute for Biology of Ageing (Cologne). Unless otherwise stated, the functions t.test() and wilcox.test() in R have been used to evaluate statistical significance. To generate a pipeline for data processing we used Snakemake^87^. Figure style was modified using Inkscape version 0.92.4. For circular visualization of genomic data we used Circos^64^.

### Inference of demographic population history with individual resequencing data

To infer the demographic history, we performed whole genome re-sequencing of single individuals from all populations resulting in mean genome coverage between 13-21x (**Table S2**). Demographic history was inferred from single individual sequencing data using Pairwise Sequential Markovian Coalescence (PSMC’ mode from MSMC2^25^). Re-sequencing of single individuals was performed with the DNA of single individuals extracted for the pooled sequencing for each examined population. The Illumina short-insert library was constructed based on a published protocol^88^. Extracted DNA (500ng) was digested with fragmentase (New England Biolabs) for 20min at 37°C, followed by end-repair and A-tailing (1.0µl NEB End-repair buffer, 0.5µl Klenow fragment, 0.5µl Taq.Polymerase, 0.2µl T4 polynucleotide kinase, 10µl reaction volume, 30min at 25°C, 30min at 75°C) and adapter ligation (NEB Quick ligase buffer 12.5µl, Quick ligase 0.5µl, 1µl adapter P1 (D50X), 1µl adapter P2 (universal), 5µM each; 20min at 20°C, 25µl reaction volume). Next, ligation mix was diluted to 50µl and used 0.583:1 volume of home-brewed SPRI beads (SPRI binding buffer: 2.5M NaCL, 20mM PEG 8000, 10mM Tric-HCL, 1mM EDTA, oh=8, 1mL TE-washed SpeedMag beads GE Healthcare, 65152105050250 per 100mL buffer) for purification. The ligation products were amplified with 9 PCR cycles using KAPA Hifi kit (Roche, P5 universal primer and P7 indexed primer D7XX). The samples were pooled and sequenced on Hiseq X. Raw sequencing reads were trimmed using Trimmomatic-0.32 (ILLUMINACLIP:illumina-adaptors.fa:3:7:7:1:true, LEADING:20, TRAILING:20, SLIDINGWINDOW:4:20, MINLEN:50)^72^. Data files were inspected with FastQC v0.11.22. Trimmed reads were subsequently mapped to the reference genome with BWA-MEM (version 0.7.12). The SAM output was converted into BAM format, sorted, and indexed via SAMTOOLS v1.3.1^61^. Filtering and realignment was conducted with PICARD v1.119 and GATK^60^. Briefly, the reads were relabeled, sorted, and indexed with AddOrReplaceReadGroups. Duplicated reads were marked with the PICARD feature MarkDuplicates and reads were realigned with first creating a target list with RealignerTargetCreator, second by IndelRealigner from the GATK suite. Resulting reads were again sorted and indexed with SAMTOOLS. Next, the guidance for PSMC’ (https://github.com/stschiff/msmc/blob/master/guide.md) was followed; VCF-files and masked files were generated with the *bamCaller.py* script (MSMC-tools package). This step requires the chromosome coverage information to mask regions with too low or too high coverage. As recommended in the guidelines, the average coverage per chromosome was calculated using SAMTOOLS. In addition, this step was performed using a coverage threshold of 18 as recommended by Nadachowska-Brzyska *et al*.^89^. Final input data was generated using the *generate_multihetsep.py* script (MSMC-tools package). Subsequently, for each sample PSMC’ was run independently. Bootstrapping was performed for 30 samples per individual and input files were generated with the *multihetsep_bootstrap.py* script (MSMCtools package).

#### Analysis of differential expressed genes with age

We downloaded the previously published RNaseq data from a longitudinal study of *Nothobranchius furzeri*^*18*^. The data set contains five time points (5w, 12w, 20w, 27w, 39w) in three different tissues (liver, brain, skin). The raw reads were mapped to the NFZ v2.0 reference genome and subsequently counted using STAR (version 2.6.0.c)^90^ and FeatureCounts (version 1.6.2)^91^. We performed statistical analysis of differential expression with age using DESeq2^92^ and age as factor. Genes are classified as upregulated in young (log(FoldChange) < 0, adjusted p < 0.01), upregulated in old (log(FoldChange) > 0,adjusted p < 0.01).

## Acknowledgments

We would like to thank Patience and Edson Gandiwa for their administrative support, Tamuka Nhiwatiwa for helping with logistics and samples handling; Evious Mpofu, Hugo and Elsabe van der Westhuizen and all the rangers of the Gonarezhou National Park for their support in the field. We are thankful to Zimbabwe National Parks for allowing our team to conduct research in the Gonarezhou National Park; Itamar Harel, Matej Polacik and Radim Blazek for hands-on contribution with the field work. We further thank all members of the Valenzano lab for their continuous scientific input and support. The Czech Science Foundation provided financial support to MR for sampling Mozambican populations (19-01789S). This project was funded by the Max Planck Institute for Biology of Ageing, the Max Planck Society and the CECAD at the University of Cologne.

---

**Figure S1.**
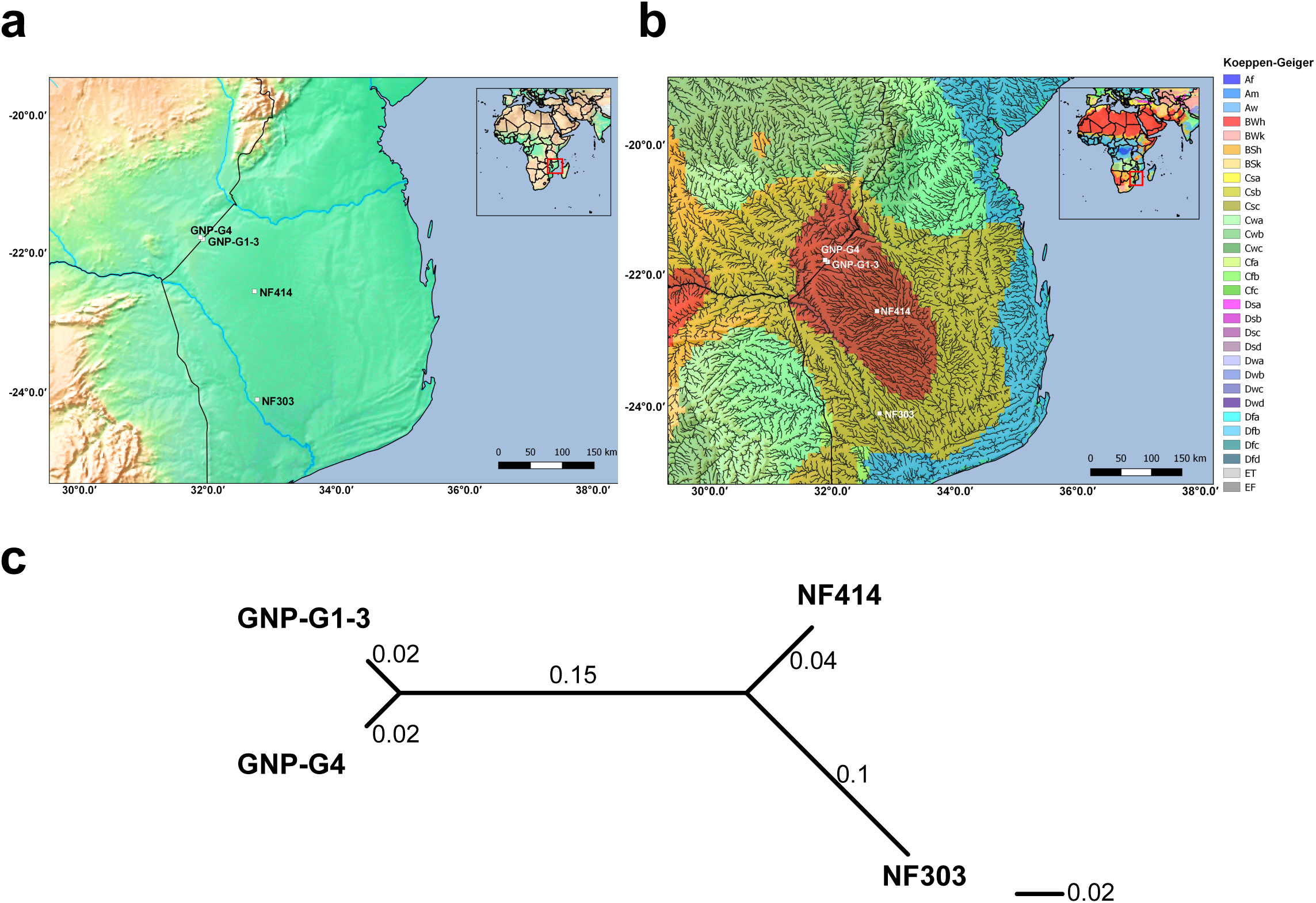
Altitude, climate classification and genetic differentiation of studied samples. a) Altitude elevation map with studied samples. b) Climate classification based on Koeppen-Geiger index combined with a high resolution river map. c) Unrooted neighbor joining tree based on pairwise genetic differentiation (F_ST_) between all sample sites.

**Figure S2.**
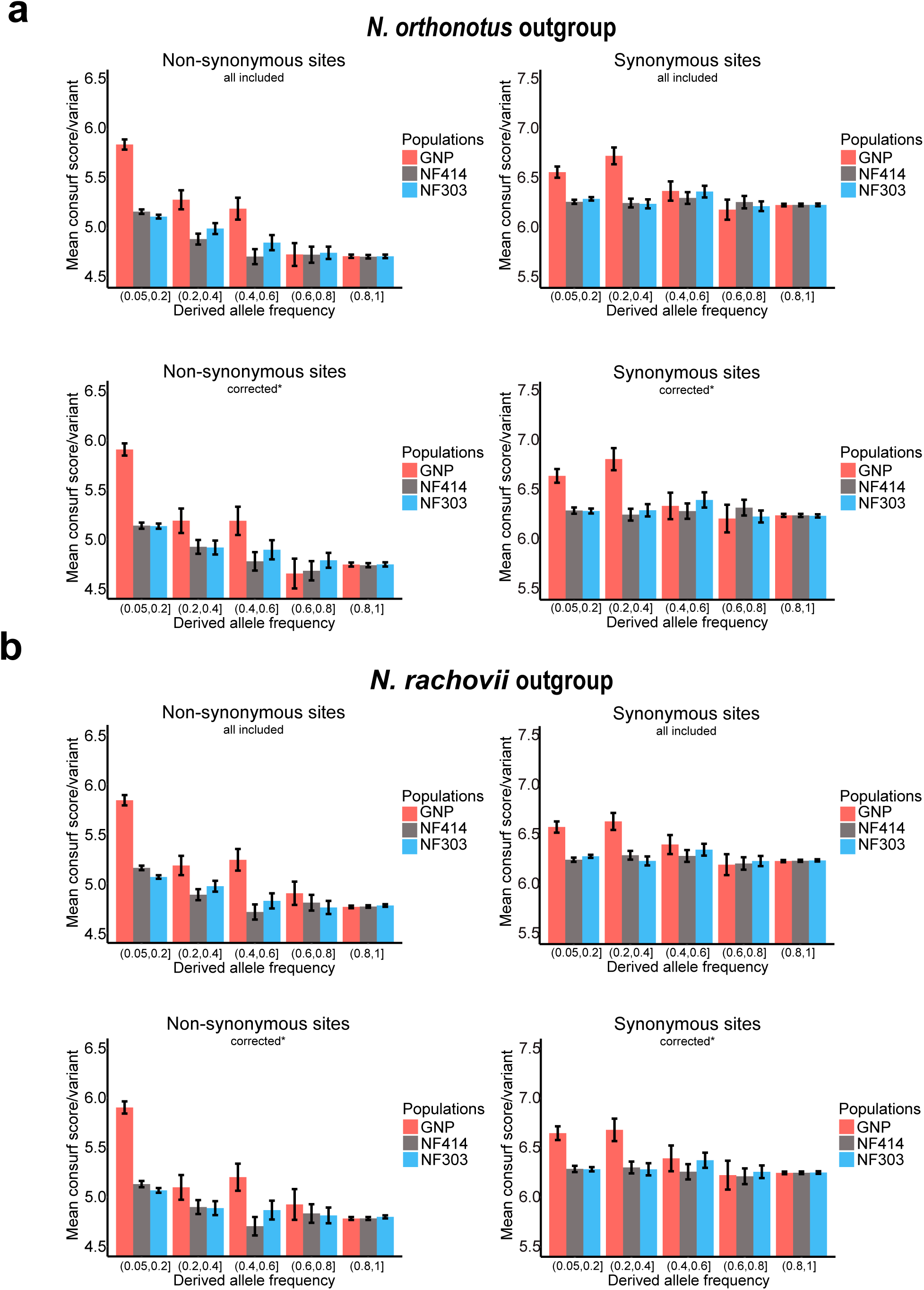
Mean Consurf score per variant based on derived frequency bins. **a)** Mean Consurf score based on derived frequency bins in non-synonymous (left) and synonymous (right) sites using *Nothobranchius orthonotus* as outgroup, including all available sites (upper panel) or only sites corrected for CMD, CpG hypermutation and highly detrimental effect based on SnpEFF analysis (lower panel). **b)** Mean Consurf score based on derived frequency bins in non-synonymous (left) and synonymous (right) sites using *Nothobranchius rachovii* as outgroup, including all available sites (upper panel) or only sites corrected for CMD, CpG hypermutation and highly detrimental effect based on SnpEFF analysis (lower panel). Mean consurf scores per variant are shown with SEM.

